# Learning from B Cell Evolution: Adaptive Multi-Expert Diffusion for Antibody Design via Online Optimization

**DOI:** 10.1101/2025.08.02.668313

**Authors:** Hanqi Feng, Peng Qiu, Meng-Chun Zhang, Yiran Tao, You Fan, Jingtao Xu, Barnabas Poczos

**Affiliations:** School of Computer Science, Carnegie Mellon University; Department of Biostatistics, University of Pittsburgh; Department of Mathematics, King’s College London; Department of Statistics, London School of Economics and Political Science

## Abstract

Recent advances in diffusion models have shown remarkable potential for antibody design, yet existing approaches apply uniform generation strategies that cannot adapt to each antigen’s unique requirements. Inspired by B cell affinity maturation—where antibodies evolve through multi-objective optimization balancing affinity, stability, and self-avoidance—we propose the first biologically-motivated framework that leverages physics-based domain knowledge within an online meta-learning system. Our method employs multiple specialized experts (van der Waals, molecular recognition, energy balance, and interface geometry) whose parameters evolve during generation based on iterative feedback, mimicking natural antibody refinement cycles. Instead of fixed protocols, this adaptive guidance discovers personalized optimization strategies for each target. Our experiments demonstrate that this approach: (1) discovers optimal SE(3)-equivariant guidance strategies for different antigen classes without pre-training, preserving molecular symmetries throughout optimization; (2) significantly enhances hotspot coverage and interface quality through target-specific adaptation, achieving balanced multi-objective optimization characteristic of therapeutic antibodies; (3) establishes a paradigm for iterative refinement where each antibody-antigen system learns its unique optimization profile through online evaluation; (4) generalizes effectively across diverse design challenges, from small epitopes to large protein interfaces, enabling precision-focused campaigns for individual targets.

## Introduction

Computational antibody design remains a fundamental challenge in therapeutic development, requiring simultaneous optimization of hotspot coverage, structural stability, and binding interface quality (Fischman and Ofran 2018; Norman et al. 2020). While recent diffusion-based methods like RFdiffusion (Watson et al. 2023) and RFAntibody (Luo et al. 2022) show promise for generating novel protein structures, they lack mechanisms to incorporate real-time constraints during generation, often producing designs that fail to meet multiple competing objectives (Eguchi et al. 2022).

Current approaches face three fundamental limitations: they generate structures through diffusion without target-specific guidance (Watson et al. 2023; Trippe et al. 2023), (2) they cannot balance multiple objectives during generation, requiring inefficient post-hoc filtering (Jin et al. 2022; Shuai, Ruffolo, and Gray 2021), and (3) they either ignore physical constraints or require extensive labeled data to train property predictors (Dauparas et al. 2022; Hsu et al. 2022). These limitations significantly reduce their effectiveness for therapeutic antibody design, where each target presents unique challenges (Akbar et al. 2022; Mason et al. 2021).

We present an adaptive guidance framework that addresses these limitations by introducing physics-aware constraints directly into the SE(3)-equivariant diffusion process (Hoogeboom et al. 2022; Corso et al. 2023). Our contributions include:

- **Multi-expert guidance system**: We develop specialized guidance modules for van der Waals interactions (Jumper et al. 2021), molecular recognition (Gainza et al. 2020), energy balance (Ingraham et al. 2019), and interface geometry (Schneider et al. 2022), each providing targeted gradients during diffusion while maintaining equivariance (Satorras, Hoogeboom, and Welling 2021).
- **Novel expert routing**: A dynamic routing system activates appropriate experts based on real-time structural metrics and diffusion timestep, ensuring optimal constraint application throughout generation (Xue et al. 2015; Olympiou et al. 2022).
- **Online parameter adaptation**: Using Bayesian optimization with Gaussian processes, our framework learns optimal guidance strategies for each antigen-antibody pair through iterative batch evaluation, automatically discovering target-specific temporal profiles without requiring pre-training (Raybould et al. 2019; Shanehsazzadeh et al. 2023).

Experiments across diverse targets demonstrate substantial improvements: 7% reduction in CDR-H3 RMSD, 9% increase in hotspot coverage, 12% better interface pAE, and 5% higher shape complementarity, with enhanced metrics across all evaluation criteria. This balanced optimization addresses the critical “weakest link” problem (Raybould et al. 2019) in antibody design. By combining physics-based guidance with online learning, our work opens new directions for adaptive approaches in biomolecular design (Ruffolo et al. 2023; Evans et al. 2022).

## Related Work

### Computational Antibody Design

The field has evolved from physics-based approaches using Rosetta (Kaufmann et al. 2010) and MODELLER (Webb and Sali 2016) to deep learning methods inspired by AlphaFold2 (Jumper et al. 2021). DeepAb (Ruffolo, Sulam, and Gray 2022) pioneered deep learning for antibody structure prediction, while ABlooper (Abanades et al. 2022) focused on CDR-H3 modeling.

Recent generative approaches shifted to de novo design. DiffAb (Luo et al. 2022) introduced diffusion models for joint sequence-structure generation but operates in internal coordinates, limiting inter-chain modeling. RFAntibody (Adolf-Bryfogle, Toth, and Bahl 2024) addressed this using SE(3)-equivariant backbone generation in Cartesian space. However, current methods lack both physicochemical constraint enforcement and target-specific adaptation during generation. This results in high-throughput campaigns producing substantial fractions of structurally unreasonable or experimentally nonviable designs (Shen et al. 2024). Additional related work is discussed in Appendix A due to space limitations.

## Preliminaries

### SE(3)-Equivariant Diffusion Models

Protein structure modeling requires respecting 3D spatial symmetries. RFdiffusion (Watson et al. 2023) combines SE(3)-equivariant networks with diffusion models, performing diffusion directly on backbone coordinates while maintaining rotational and translational equivariance.

#### Frame Representations and Forward Diffusion Process

Each residue *i* is represented by a rigid body frame **T**_*i*_ = (**R**_*i*_, **t**_*i*_) *∈* SE(3), where **R**_*i*_ *∈* SO(3) is a rotation matrix and **t**_*i*_ *∈* R^3^ is a translation vector. For *N* residues, the complete structure is **T** =*{* **T**_1_, …, **T**_*N*_ *}*.

The forward diffusion process adds noise over time steps *t ∈ {*0, 1, …, *T}*. Rotations follow the IGSO3 distribution, *q*(**R**_*t*_|**R**_0_) = IGSO3(**R**_*t*_; **R**_0_, *σ*_*t*_) with density:

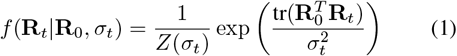

where *σ*_*t*_ is the noise level at timestep *t*, controlling the variance of the distribution(Yim et al. 2023). The derivation of the score function and its properties on the SO(3) manifold are detailed in Appendix B. Translations follow Gaussian diffusion with center-of-mass constraint:

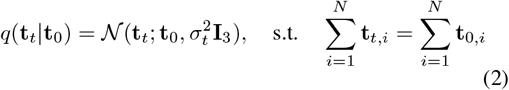

#### Reverse Diffusion and Network Architecture

The reverse process learns to denoise the forward diffusion process by estimating clean structure **T**_0_ from noisy **T**_*t*_ (Ho, Jain, and Abbeel 2020):

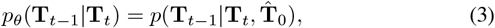

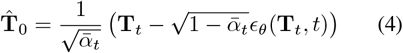

Where 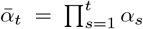 is the cumulative product of noise schedule parameters, and *ϵ*_*θ*_(**T**_*t*_, *t*) is the neural network with parameters *θ* that predicts the noise added at timestep *t*.

### RFAntibody Pipeline

RFAntibody (Adolf-Bryfogle et al. 2024) employs a three-stage pipeline for antibody design: (1) RFdiffusion generates diverse antibody backbones conditioned on the target antigen, (2) ProteinMPNN designs sequences compatible with these structures, and (3) RoseTTAFold2 validates the designs by predicting their folded structures. This decoupled approach allows each model to focus on its specialized task—structure generation, sequence design, and validation respectively. The pipeline filters designs based on structural quality metrics, retaining only those most likely to succeed experimentally.

## Problem Formulation and Motivation

While state-of-the-art generative models excels at generating diverse protein structures, it faces a critical challenge in antibody design: maintaining balanced performance across multiple essential metrics. In antibody engineering, success requires simultaneous optimization of structural accuracy (CDR-H3 RMSD), prediction confidence (pAE, ipAE), interface quality (shape complementarity, buried surface area), and biophysical properties (VDW interactions, stability) (Raybould et al. 2019; Norman et al. 2020; Choi et al. 2018).

The fundamental issue is that RFdiffusion’s purely data-driven approach lacks mechanisms to enforce multiobjective balance. This leads to the “weakest link” problem—a single failing metric renders the entire design experimentally nonviable (Raybould et al. 2019; Choi et al. 2018), regardless of excellence in other areas. In high-throughput campaigns, even if 90% of metrics are excellent, failure in the remaining 10% eliminates the design from experimental consideration.

Our work addresses this challenge by developing an adaptive guidance system that steers the diffusion process toward regions where all objectives are simultaneously satisfied. Unlike post-hoc filtering approaches that discard failed designs, our method proactively guides generation to maintain metric balance throughout the process, significantly improving the yield of experimentally viable candidates per computational cycle (Figure 1).

**Figure 1:**
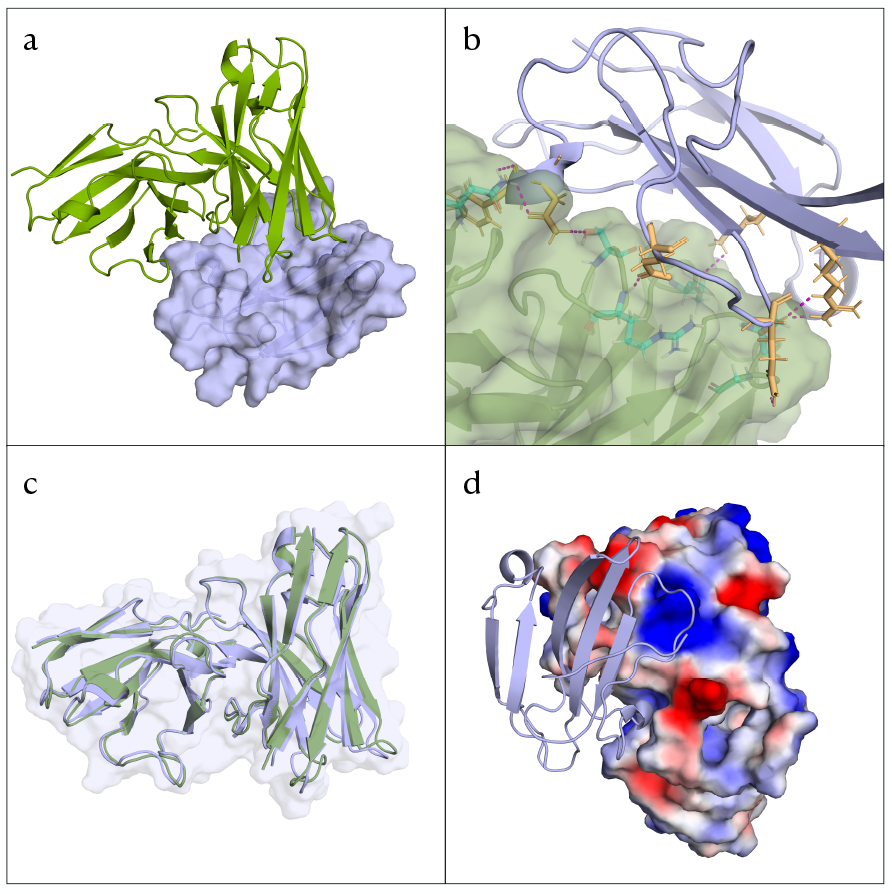
High-quality antibody design showcasing the effectiveness of our adaptive guidance. (a) Overview of the antibody-antigen complex. (b) Detailed view of the binding interface demonstrating enhanced contact density achieved through guided optimization. (c) Structural alignment showing exceptional CDR-H3 accuracy (RMSD = 0.63 Å). (d) Electrostatic surface representation illustrating improved charge complementarity resulting from physics-aware guidance.

## Method

### Theoretical Foundation

#### Biophysical Principles of Antibody-Antigen Recognition

Our multi-expert system is grounded in the fundamental physics of molecular recognition. Antibody-antigen binding emerges from a delicate balance of forces:

#### Energetic Landscape

The binding free energy (Δ*G*_bind_) can be decomposed as (Gilson and Zhou 2007):

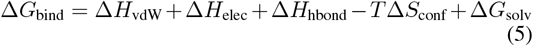

where each term corresponds to van der Waals, electrostatic, hydrogen bonding, conformational entropy, and solvation contributions respectively. Our expert modules directly optimize these physical components.

#### Shape Complementarity Principle

Following the lockand-key model extended by induced fit theory (Fischer 1894; Koshland 1958), optimal binding requires:

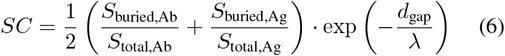

where Ab and Ag denote antibody and antigen respectively, *S*_buried_ represents buried surface area upon complex formation, *S*_total_ is the total surface area of each protein, and *d*_gap_ captures interface gaps penalized by decay length *λ*.

#### Information-Theoretic Justification for Adaptive Guidance

Our adaptive expert routing is motivated by information theory. At timestep *t*, the mutual information between the noisy structure **x**_*t*_ and the target structure **x**_0_ is (Austin et al. 2021):

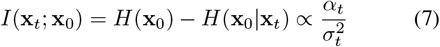

where *α*_*t*_ is the signal retention coefficient at timestep *t*. The derivation of this relationship is provided in Appendix C.

This suggests that different structural features become identifiable at different noise levels, justifying our adaptive expert routing based on the signal-to-noise ratio at each timestep. The connection between SNR and feature visibility is detailed in Appendix D.

### Overview

We present an adaptive physics-guided diffusion framework for antibody design that integrates multi-expert guidance with online learning. Our approach enhances the standard SE(3) equivariant diffusion process (Hoogeboom et al. 2022; Watson et al. 2023) by incorporating domain-specific physical constraints through a dynamically weighted ensemble of expert modules. Figure 2 presents our adaptive framework. The system combines physics-based expert guidance (left) with the antibody design pipeline, where guided diffusion generates structures, ProteinMPNN designs sequences, and RoseTTAFold2 validates results. A Bayesian optimization feedback loop (bottom) continuously learns optimal guidance parameters from evaluation metrics.

**Figure 2:**
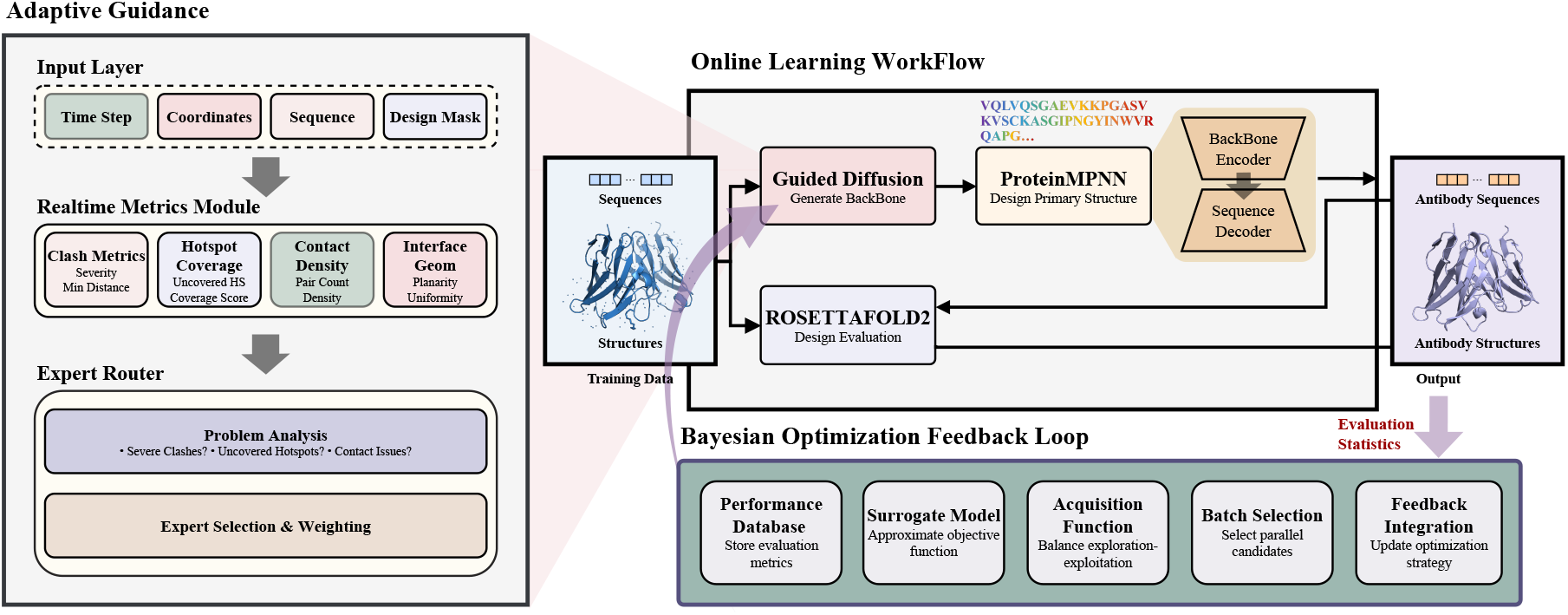
Adaptive multi-expert diffusion framework with online learning for antibody design. Due to space constraints, detailed descriptions of each module and their input/output dimensions are provided in Appendix F.

### Physics-Guided SE(3) Diffusion

Our framework modifies the standard reverse diffusion process by incorporating physics-based guidance gradients. The reverse step is formulated as:

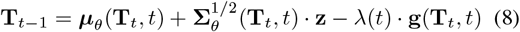

where ***µ***_*θ*_ and **Σ**_*θ*_ are the predicted mean and covariance from the denoising network, **z ∼** N (0, **I**), and **g**(**X**_*t*_, *t*) represents the combined guidance gradient:

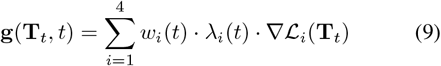

where *w*_*i*_(*t*) are adaptive weights for each expert and *∇L*_*i*_ are gradients of expert-specific physics-based loss functions.

The guidance strength for each expert *i* at timestep *t* is determined as:

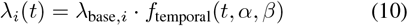

where *λ*_base,*i*_ is the base strength for expert *i*, and *f*_temporal_(*t, α, β*) controls when each expert is most active during denoising. The shape parameters *α* and *β* are learned via Bayesian optimization to discover optimal temporal profiles—early-peaking (*α < β*) for global structure experts, late-peaking (*α > β*) for atomic refinement:

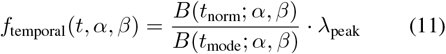

where *t*_norm_ = (*t −* 1)*/*(*T −* 1) normalizes the timestep to [0, 1], and *B*(·; *α, β*) is the Beta density function. The shape parameters *α* and *β* are learned via Bayesian optimization to discover optimal activation timing for each target. Detailed profiles are shown in Appendix E.

### Multi-Expert Guidance System

We employ four specialized expert modules, each addressing critical aspects of antibody-antigen interactions:

#### VDW Balance Expert

This expert prevents steric clashes while maintaining favorable van der Waals interactions. The loss function penalizes atomic overlaps:

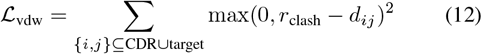

where *r*_clash_ is the clash threshold and *d*_*ij*_ = ∥ **x**_*i*_ *−* **x**_*j*_ ∥is the distance between backbone atoms. The gradient is:

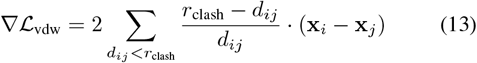

#### Molecular Recognition Expert

This expert ensures proper CDR coverage of epitope hotspots. For each hotspot *h ∈ H*, we encourage proximity to CDR residues(see Appendix G for hotspot definition and selection methodology):

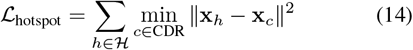

The gradient attracts the nearest CDR residue *c*^*∗*^ toward uncovered hotspots:

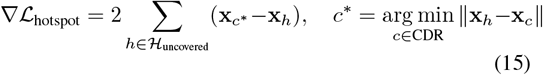

where *H*_uncovered_ denotes inadequately covered hotspots.

#### Energy Balance Expert

This expert maintains optimal contact density at the binding interface. The loss function is defined as:

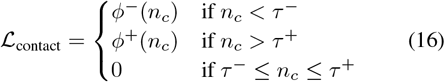

where *n*_*c*_ denotes the number of inter-molecular contacts within threshold *d*_*c*_, [*τ*^*−*^, *τ* ^+^] defines the target range, and *ϕ*^*−*^(·), *ϕ*^+^(·) are penalty functions for sparse and dense packing, respectively. The gradient modulates CDR positioning to optimize interface contacts.

#### Interface Quality Expert

This expert optimizes the geometric properties of the binding interface through multiple physically motivated objectives:

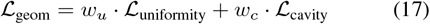

where *w*_*u*_ and *w*_*c*_ are weighting coefficients that balance the contribution of each geometric criterion.

#### Distance Uniformity

The uniformity of residue spacing is evaluated to ensure well-distributed interface contacts:

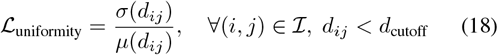

where *d*_*ij*_ represents pairwise distances between interface residues *I*, and *σ*(·), *µ*(·) denote standard deviation and mean, respectively. This metric prevents local clustering or sparse regions at the interface.

#### Cavity Detection

The cavity penalty identifies isolated residues lacking sufficient neighbors:

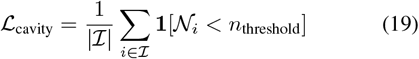

where *N* _*i*_ counts neighbors within radius *r*_neighbor_, and **1**[·] is the indicator function that equals 1 if the condition is true and 0 otherwise. This penalty prevents the formation of hydrophobic cavities and ensures a well-packed interface.

### Adaptive Expert Routing

Our routing mechanism dynamically activates experts based on real-time structural analysis, ensuring that computational resources focus on the most critical problems while maintaining SE(3)-equivariance. The system computes problem severity scores:

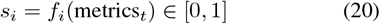

where *f*_*i*_ evaluates current structural metrics to determine the severity of problems relevant to expert *i* (detailed in the Appendix H).

The activation is problem-driven rather than time-dependent. For instance:

- High clash density (*d*_min_ *< r*_clash_) triggers VDW expert activation
- Low hotspot coverage activates molecular recognition guidance
- Suboptimal contact density engages energy balance optimization
- Poor interface geometry activates geometric refinement Experts are weighted proportionally to problem severity:

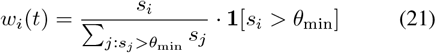

This adaptive weighting ensures that only experts addressing detected problems are activated, preventing unnecessary or conflicting guidance.

Critically, SE(3)-equivariance is preserved because all expert gradients are computed from invariant geometric features. Each expert’s gradient satisfies:

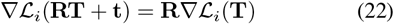

for any rotation **R** *∈* SO(3) and translation **t** *∈* ℝ^3^. This is achieved by basing all computations on pairwise distances, relative orientations, and other invariant descriptors rather than absolute coordinates. The weighted combination of equivariant gradients maintains equivariance: 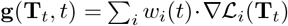 remains equivariant since weights *w*_*i*_(*t*) are scalar functions of invariant metrics.

### Online Parameter Learning

We employ Bayesian optimization with Gaussian processes (GP) to adaptively learn the optimal Beta distribution parameters for each antibody-antigen system post-generation. Our approach models a two-dimensional parameter space ***θ*** = (*α, β*) *∈ Θ*, where *α* and *β* are the shape parameters controlling the temporal modulation profile. The optimization process maintains a probabilistic surrogate model of the objective function using a Gaussian process:

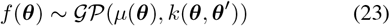

where *µ*(***θ***) represents the predicted mean performance and *k*(***θ, θ***^*′*^) is the covariance function. We employ a Matérn kernel with automatic noise estimation:

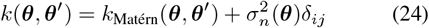

After each design evaluation, the GP posterior is updated using the observed loss:

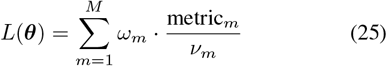

where the metrics include CDR-H3 backbone RMSD, predicted aligned error (pAE), and interaction pAE (ipAE), each normalized by appropriate constants *ν*_*m*_.

The next parameter configuration is selected by maximizing the Expected Improvement (EI) acquisition function:

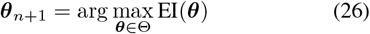

To handle stochastic evaluations, we aggregate results from similar parameters to reduce noise and periodically reevaluate promising configurations. This approach automatically discovers optimal Beta parameters for each target without manual tuning. Due to space constraints, convergence analysis and computational requirements are detailed in Appendix I.

## Experiments

### Experimental Setup

#### Dataset and Targets

We evaluated our framework on six antibody-antigen pairs: IL17A (PDB: 6PPG), ACVR2B (PDB: 5NGV), FXI (PDB: 6HHC), TSLP (PDB: 5J13), IL36R (PDB: 6U6U), and TNFRSF9 (PDB: 6A3W). Prior work including RFAntibody (Bennett et al. 2024) typically evaluated four targets; we expanded this to six therapeutically relevant targets spanning inflammation, thrombosis, and cancer applications (17-40 kDa). This extended benchmark from IgDesign (Shanehsazzadeh et al. 2023) includes 1,243 SPR-validated designs across diverse binding modes. We annotated 5 hotspot residues per target and generated 1,000 designs each, following the field’s established protocol for rigorous benchmarking of antibody design methods.

#### Baseline Methods

Most recent methods are unavailable due to lack of publicly accessible code. We selected: **DiffAb** (Luo et al. 2022): torsion-space CDR generation; **RFAntibody** (Bennett et al. 2024): the current state-of-theart method without physics guidance, widely recognized for its exceptional performance across diverse antibody design tasks. For fair comparison, all methods used identical inputs (target structure, hotspot annotations, and Trastuzumab framework).

#### Implementation Details

All experiments used 4 NVIDIA RTX A6000 GPUs with the following settings in Table 1.

**Table 1:** Implementation details of our adaptive physicsguided framework.

| Parameter | Value | Parameter | Value |
| --- | --- | --- | --- |
| *Diffusion & Guidance* |  | *Beta Distribution* |  |
| Timesteps $T$ | 50 | $\alpha$ range | [0.5, 10.0] |
| Expert modules | 4 | $\beta$ range | [0.5, 10.0] |
| VDW threshold | 2.8 Å | Peak factor $\lambda_{\text{peak}}$ | 5.0 |
| Gradient clip | 2.0 | Initial $(\alpha, \beta)$ | (2.0, 2.0) |
| Distance cutoff | 8.0 Å |  |  |
| *Expert Activation* |  | *Online Learning* |  |
| Min threshold $\theta_{\min}$ | 0.1 | Kernel | Matérn-5/2 |
| VDW strength | 0.5 | Length scale | 2.0 |
| Recognition strength | 2.0 | GP noise | 0.3 |
| Activation | Adaptive | Acquisition $\xi$ | 0.01 |
| *Evaluation Metrics* |  | *Implementation* |  |
| RMSD norm $\nu_1$ | 1.5 Å | Framework | PyTorch |
| pAE norm $\nu_2$ | 7.0 | GPU memory | 48GB |
| ipAE norm $\nu_3$ | 10.0 | CDR design | All 6 |

### Evaluation Metrics

We employ a comprehensive set of metrics to evaluate antibody design quality across structural accuracy, binding interface characteristics, and biophysical properties.

#### CDR-H3 RMSD

We measure the backbone root-meansquare deviation of the critical CDR-H3 loop after structural alignment of framework regions. CDR-H3 is the most diverse and challenging loop to model, directly correlating with binding specificity.

#### Predicted Alignment Error (pAE)

Extracted from AlphaFold2-Multimer confidence predictions, pAE estimates the expected position error between residue pairs. We report both mean pAE across the entire complex and mean interaction pAE (ipAE) specifically for residue pairs across the antibody-antigen interface. Lower values indicate higher confidence in the predicted structure.

#### Predicted Local Distance Difference Test (pLDDT)

Per-residue confidence metric. We average pLDDT across CDR residues for local structural quality assessment.

#### Shape Complementarity (SC)

Quantifies geometric fit between antibody-antigen surfaces, with higher values indicating better complementarity.

#### Buried Surface Area (BSA)

Solvent-accessible surface area buried upon complex formation. Larger interfaces correlate with stronger, more specific interactions.

#### Hotspot Coverage

Percentage of critical epitope residues forming close contacts with CDR residues.

#### CDR Participation

Fraction of CDR residues involved in antigen binding. Higher participation indicates more distributed binding.

#### Energetic and Biophysical Metrics

##### Van der Waals Energy

We compute packing interactions and steric clashes using physics-based energy functions. Favorable scores indicate well-packed interfaces without significant clashes.

##### Overall Success Criteria

Successful designs must satisfy: CDR-H3 RMSD *<* 3.0 Å, mean pAE *<* 10.0, and mean interaction pAE (ipAE) *<* 10.0. This multi-metric evaluation ensures structural accuracy and binding confidence.

This multi-metric evaluation ensures designs are not only structurally accurate but also likely to bind with high affinity and specificity.

### Main Results

Table 2 presents comprehensive evaluation across all targets. Our method achieved improvements in most metrics while maintaining a more balanced performance profile compared to both baselines, effectively addressing the critical “weakest link” problem in antibody design where poor performance in any single metric can compromise experimental viability. As shown in Figure 3 and Figure 4, our adaptive guidance consistently improves performance across all key metrics (CDR-H3 RMSD, pAE, ipAE) for multiple targets, with notably reduced variance in the distributions.

**Table 2:**
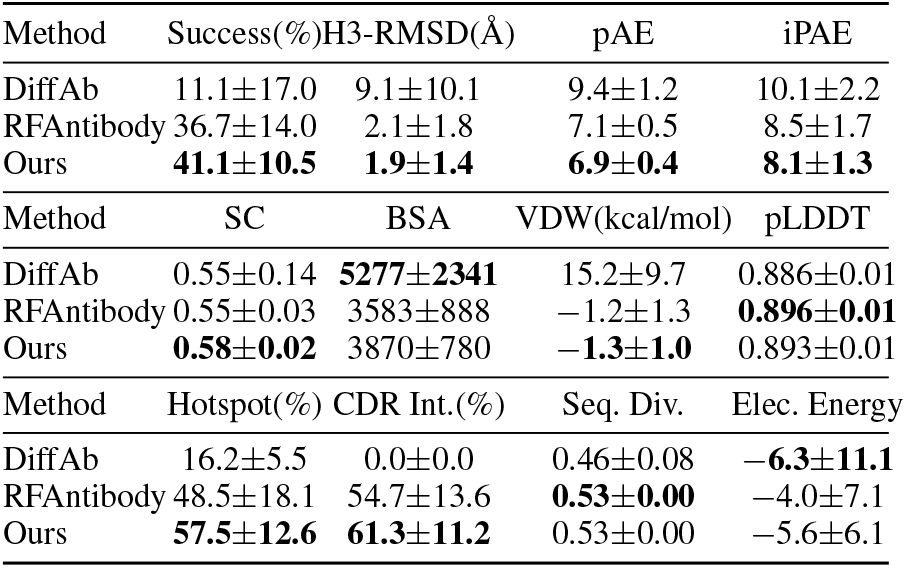
Overall performance comparison across six targets highlighting balanced optimization.

| Method | Success(%) | H3-RMSD(Å) | pAE | iPAE |
| --- | --- | --- | --- | --- |
| DiffAb | 11.1±17.0 | 9.1±10.1 | 9.4±1.2 | 10.1±2.2 |
| RFAntibody | 36.7±14.0 | 2.1±1.8 | 7.1±0.5 | 8.5±1.7 |
| Ours | **41.1±10.5** | **1.9±1.4** | **6.9±0.4** | **8.1±1.3** |

| Method | SC | BSA | VDW(kcal/mol) | pLDDT |
| --- | --- | --- | --- | --- |
| DiffAb | 0.55±0.14 | **5277±2341** | 15.2±9.7 | 0.886±0.01 |
| RFAntibody | 0.55±0.03 | 3583±888 | -1.2±1.3 | **0.896±0.01** |
| Ours | **0.58±0.02** | 3870±780 | **-1.3±1.0** | 0.893±0.01 |

| Method | Hotspot(%) | CDR Int.(%) | Seq. Div. | Elec. Energy |
| --- | --- | --- | --- | --- |
| DiffAb | 16.2±5.5 | 0.0±0.0 | 0.46±0.08 | -6.3±11.1 |
| RFAntibody | 48.5±18.1 | 54.7±13.6 | **0.53±0.00** | -4.0±7.1 |
| Ours | **57.5±12.6** | **61.3±11.2** | 0.53±0.00 | -5.6±6.1 |

**Figure 3:**
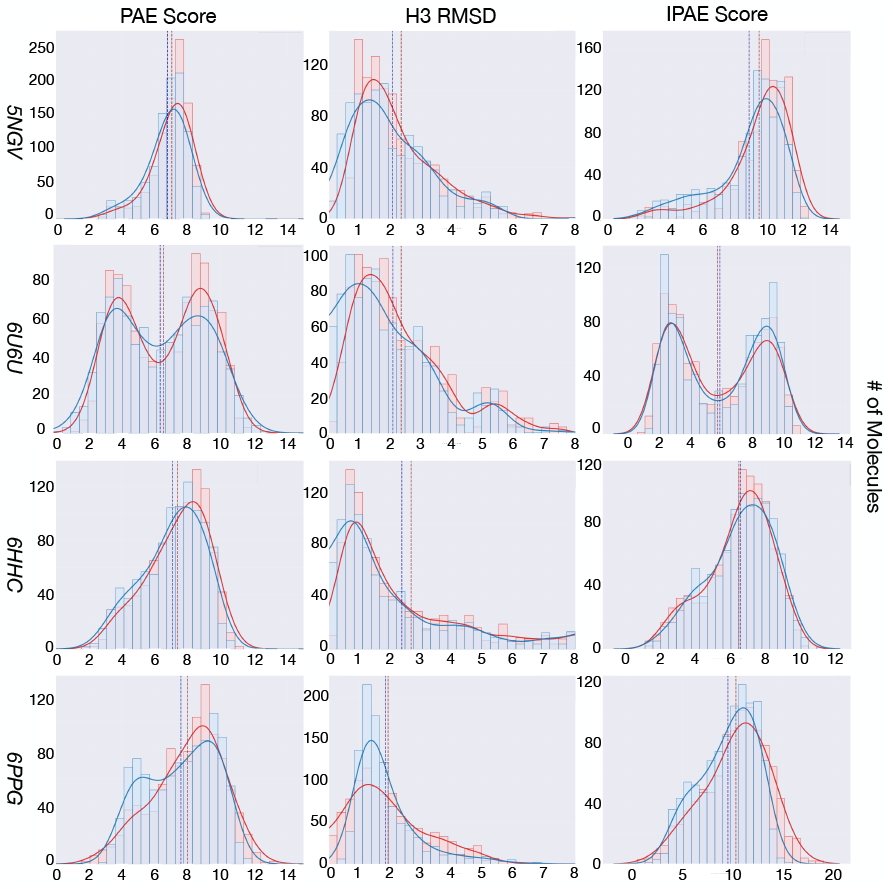
Histogram distributions of evaluation metrics (pAE, CDR-H3 RMSD, ipAE) for baseline (red) and adaptive guidance (blue) methods across four antibody targets.

**Figure 4:**
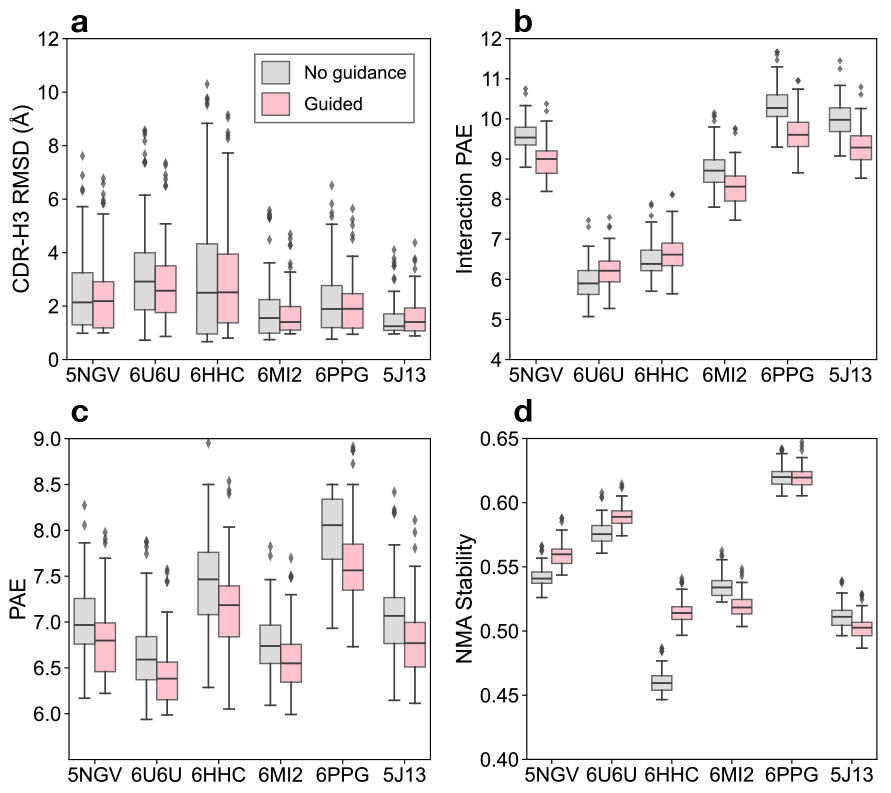
Adaptive guidance improves antibody design quality across multiple metrics. Box plots compare baseline RFAntibody (gray) with our guided approach (pink) on six targets for (a) CDR-H3 RMSD, (b) interaction pAE, (c) mean pAE, and (d) NMA stability.

#### Online Learning Impact

To evaluate the contribution of our online learning framework, we conducted ablation studies comparing three configurations: (1) the full system with both multi-expert guidance and online learning, (2) multi-expert guidance with fixed parameters (no online learning), and (3) the baseline RFAntibody without any guidance.

The results reveal the complementary benefits of multi-expert guidance and online learning:

#### Multi-Expert Guidance Alone

Adding physics-based guidance to RFAntibody improves the success rate from 36.7% to 38.9% (Table 3). While all metrics show improvement, the gains are modest and the standard deviations remain relatively high. This suggests that fixed guidance parameters, though helpful, cannot adapt to the diverse characteristics of different antibody-antigen systems.

**Table 3:** Ablation study demonstrating the importance of online learning.

| Config | Success | H3-RMSD | pAE | iPAE |
| --- | --- | --- | --- | --- |
| RFdiffusion | 36.7±14.0 | 2.11±1.81 | 7.13±0.46 | 8.49±1.71 |
| + Expert | 38.9±12.5 | 2.01±1.58 | 6.95±0.42 | 8.28±1.65 |
| **+ Full** | **41.1±10.5** | **1.94±1.36** | **6.85±0.39** | **8.12±1.31** |

| Config | SC | BSA | VDW | pLDDT |
| --- | --- | --- | --- | --- |
| RFdiffusion | 0.549±.026 | 3583±888 | -1.18±1.27 | 0.896±.011 |
| + Expert | 0.562±.026 | 3712±885 | -1.25±1.15 | 0.885±.014 |
| **+ Full** | **0.576±.024** | **3870±780** | **-1.33±0.95** | **0.893±.008** |

| Config | Hotspot | CDR | Seq.D | Elec.E |
| --- | --- | --- | --- | --- |
| RFdiffusion | 48.5±18.1 | 54.7±13.6 | 0.534±.004 | -3.96±7.1 |
| + Expert | 55.6±15.2 | 60.8±13.5 | 0.532±.005 | -5.31±8.1 |
| **+ Full** | **57.5±12.6** | **61.3±11.2** | **0.532±.004** | **-5.64±6.1** |

#### Online Learning Enhancement

The addition of online parameter optimization yields substantial improvements across all metrics. Most notably, the standard deviations decrease significantly, indicating more consistent and reliable design quality. This variance reduction is crucial for practical applications where predictability is as important as average performance.

#### Parameter Adaptation Analysis

Our online learning framework employs Gaussian Process-based Bayesian optimization to discover optimal Beta distribution parameters for each antibody-antigen system (Table 4).

**Table 4:** Target-specific Beta distribution parameters discovered through Bayesian optimization.

| Target Type | Beta Params |  | Key Expert Emphasis |
| --- | --- | --- | --- |
| | $\alpha$ | $\beta$ | |
| Small epitopes (IL17A, IL36R) | 3.2±0.4 | 1.8±0.3 | VDW + Recog. |
| Large interfaces (ACVR2B, TNFRSF9) | 2.1±0.3 | 3.5±0.5 | Interface + Energy |
| Buried epitopes (FXI) | 2.8±0.4 | 2.2±0.3 | Recog. + Energy |
| Flexible targets (TSLP) | 2.0±0.5 | 2.0±0.5 | All experts balanced |

## Conclusion

We presented an adaptive physics-guided framework that mimics B cell affinity maturation to optimize antibody design through online learning. This work represents the first application of adaptive guidance to diffusion-based antibody generation, demonstrating that computational design can benefit from mimicking natural immune optimization strategies.

Our key contributions include: (1) formulating antibody design as an adaptive process inspired by B cell affinity maturation, (2) pioneering online optimization for diffusion model guidance without retraining, and (3) developing a multi-expert system that achieves balanced optimization across competing objectives, mirroring natural selection pressures.

The biological significance extends beyond technical metrics. Like B cells iteratively refining antibodies through mutation and selection, our framework continuously learns from each design attempt. This parallel to immune evolution enables rapid adaptation to new antigens—critical for emerging pathogen response and personalized therapeutics. By integrating biological principles with machine learning, we bridge immunology and AI to accelerate therapeutic development.

## Limitations and Future Work

Due to space constraints, a comprehensive discussion of limitations and future research directions is provided in Appendix J.

